# Hypomorphic mutation in the large subunit of replication protein A affects mutagenesis by human APOBEC cytidine deaminases in yeast

**DOI:** 10.1101/2024.06.27.601081

**Authors:** Matthew S. Dennen, Zachary W. Kockler, Steven A. Roberts, Adam B. Burkholder, Leszek J. Klimczak, Dmitry A. Gordenin

## Abstract

Human APOBEC single-strand (ss) specific DNA and RNA cytidine deaminases change cytosines to uracils and function in antiviral innate immunity, RNA editing, and can cause hypermutation in chromosomes. The resulting uracils can be directly replicated, resulting in C to T mutations, or uracil-DNA glycosylase can convert the uracils to abasic (AP) sites which are then fixed as C to T or C to G mutations by translesion DNA polymerases. We noticed that in yeast and in human cancers, contributions of C to T and C to G mutations depends on the origin of ssDNA mutagenized by APOBECs. Since ssDNA in eukaryotic genomes readily binds to replication protein A (RPA) we asked if RPA could affect APOBEC-induced mutation spectrum in yeast. For that purpose, we expressed human APOBECs in the wild-type yeast and in strains carrying a hypomorph mutation *rfa1-t33* in the large RPA subunit. We confirmed that the *rfa1-t33* allele can facilitate mutagenesis by APOBECs. We also found that the *rfa1-t33* mutation changed the ratio of APOBEC3A-induced T to C and T to G mutations in replicating yeast to resemble a ratio observed in long-persistent ssDNA in yeast and in cancers. We present the data suggesting that RPA may shield APOBEC formed uracils in ssDNA from Ung1, thereby facilitating C to T mutagenesis through the accurate copying of uracils by replicative DNA polymerases. Unexpectedly, we also found that for uracils shielded from Ung1 by wild-type RPA the mutagenic outcome is reduced in the presence of translesion DNA polymerase zeta.

## INTRODUCTION

Humans have several APOBEC (<u>Apo</u>lipoprotein <u>B</u> mRNA-<u>E</u>diting <u>E</u>nzyme, <u>C</u>atalytic polypeptide-like) cytosine deaminases that can convert cytosines to uracils in single-stranded (ss) RNA or DNA. APOBECs operate in a physiologically important manner by editing selected mRNAs and contributing into innate immunity defense against RNA and DNA viruses (Banerjee *et al*. 2008; Harris and Dudley 2015; Lerner *et al*. 2018). Further, APOBECs can also introduce mutations into chromosomes of human malignant tumors. In fact, APOBECs are one of the most ubiquitous and prevailing causes of mutagenesis in cancers (Roberts *et al*. 2013; Mertz *et al*. 2022). Due to structural constraints, only ssDNA or RNA can be substrates for APOBEC cytidine deamination (Salter *et al*. 2016; Kouno *et al*. 2017). Short-lived stretches of ssDNA that are formed during replication, repair and in transient R-loops during transcription can be mutated by APOBECs. Alternatively, long persistent stretches of ssDNA formed by DNA end-resection in DSBs, at uncapped telomeres, or by break-induced replication can be deaminated simultaneously resulting in several cytosines being deaminated stretching over many kilobases, termed mutation clusters (Saini and Gordenin 2020). Such clusters were observed in yeast and human cell models as well as in cancer genomes. (Chan and Gordenin 2015; Sakofsky *et al*. 2019; Saini and Gordenin 2020).

Uracils formed through APOBEC cytidine deamination in ssDNA can be accurately copied by replicative DNA polymerases resulting in C to T mutations (Figure 1; steps b2→b2.1→b2.1.1). Another path of uracils-associated mutagenesis can be triggered by uracil-DNA glycosylase (Udg), which is encoded by UNG1 gene in yeast or by UNG in humans. Udg glycosylation (U-glycosylation, step b1 on Figure 1) leaves abasic (AP) site in place of an uracil. Since AP-sites are chemically unstable they may result in ssDNA breakage, rearrangements, chromosome loss or even cell death (step (c) on Figure 1). Detrimental effects of AP-sites in ssDNA can be alleviated by several ways (Boiteux and Jinks-Robertson 2013; Krokan *et al*. 2014; Saini and Gordenin 2020). If an AP-site is formed in ssDNA which can be re-annealed with undamaged complementary strand by replication fork regression or by re-winding of transiently unwound DNA within a R-loop, it will be fixed by templated error-free base excision repair (BER) without leaving any mutation trace (steps b1.2→b1.2.1 on Figure 1). Alternatively, DNA strands containing an AP-site can be copied directly by error-prone translesion synthesis (TLS) which will result in APOBEC-induced mutations (steps b1.1→b1.1.1 on Figure 1). All extensions from a base inserted across an AP-site require Pol zeta polymerase (catalytic subunit encoded by *REV3* gene in yeast). Pol zeta extension complex also includes non-catalytic subunits of replicative polymerase delta, Rev7 subunit and Rev1 (Martin and Wood 2019). The Rev1 serves as scaffold for the Pol zeta extension step as well as to perform TLS insertion of a cytosine across AP-site. Polymerase(s) inserting adenines or thymines are not yet defined ((Chan *et al*. 2012; Boiteux and Jinks-Robertson 2013; Chan *et al*. 2013; Hoopes *et al*. 2017) and references therein). Thus, U-glycosylation followed by TLS across AP-site and subsequent DNA replication (step b1.1.1 on Figure 1) can result in any of the three possible substitutions of a cytosine, C to T, C to G, and C to A as well as in a non-mutagenic outcome. Summarizing pathways illustrated on Figure 1, the spectrum of three possible APOBEC-induced substitutions of cytosines would be defined by the efficiency of U-glycosylation and by relative contribution of mutagenic TLS pathways.

**Figure 1.**
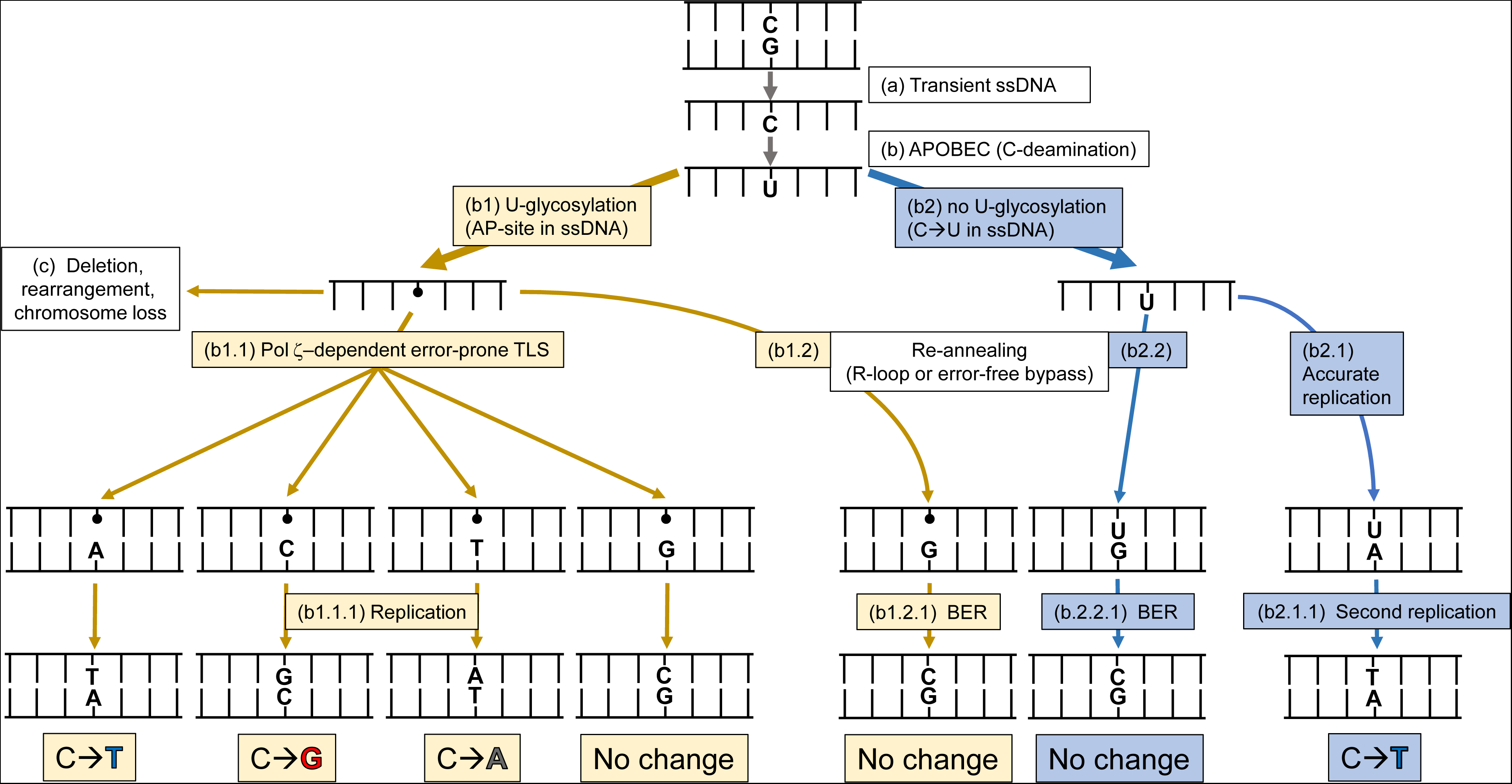
Channeling a uracil (U) created by APOBEC deamination of cytosine (C) in transient single-stranded (ss) DNA via error-prone and error-free replication and repair pathways. Pathways and steps are identified by the combined letter and legal numbering styles. **(a)** Transient ssDNA intermediates can be formed through a range of DNA replication, transcription, and repair events (see Figures 1 and 2 in (Saini and Gordenin 2020) and references therein). **(b)** ssDNA-specific APOBEC cytosine deaminase converts C to U and then is processed via sub-pathways **(b1)** and **(b2)**, which are color-coded for distinction. Sub-pathway **(b1)** starts from abasic site (AP, shown as a small ball) created by uracil-DNA glycosylase (Ung1 in yeast). AP-site can lead to DNA breakage, rearrangements to chromosome loss - box **(c)**. It can also result in base-substitutions. Another sub-pathway involves unchanged U (sub-pathway **(b2**)). Restoration of transient ssDNA containing an AP site to double-stranded (ds) form can be performed by DNA polymerase(s), including by Pol-zeta- dependent error prone translesion synthesis (TLS) **(b.1.1)**, Next round of DNA replication **(b.1.1.1)** will fix a base substitution or a non-mutant sequence. Restoration to of ssDNA to dsDNA can also occur by re-annealing (sub-pathways **(b.1.2)**) and **(b.1.2)**. Re-annealing of ssDNA can involve a complementary strand of the same DNA molecule when ssDNA was formed by transient unwinding or within the R-loop transcription intermediate. It can also involve a complimentary strand of a sister DNA molecule, if re-annealing occurs via replication fork regression. Re-annealed of AP-containing dsDNA can be repaired by base-excision repair (BER) utilizing a complimentary strand with wild type sequence as a template **(b.1.2)**, restoring wild type dsDNA sequence. Similarly, re-annealing involving U-containing ssDNA **(b.2.1)** would be a subject to BER fixing the wild type sequence in resulting dsDNA. On the contrary, two rounds of accurate replication of U-containing ssDNA **(b.2.2)** and **(b.2.2.1)** would always generate a C→T mutation.

In human cancers as well as in the yeast model systems, APOBEC mutagenesis produced mostly C to T and C to G mutations with C to A mutations occurring at low frequency barely distinguishable from non-APOBEC mutation background (Chan *et al*. 2013; Chan and Gordenin 2015; Mertz *et al*. 2022). We noticed that in UNG1 wild-type (WT) yeast the ratio of C to T and C to G mutations depended on the way ssDNA was formed (Figure 2). In yeast undergoing normal replication, APOBEC-induced C to T mutations strongly prevailed over C to G events and both rarely formed mutation clusters, i.e., were scattered over the genome. On the contrary, the spectra of APOBEC-induced clustered mutations in long persistent ssDNA formed by end-resection at uncapped telomeres or via break-induced replication (BIR) contained comparable numbers of C to G and C to T events. Mutation clusters in human cancers enriched with mutations in APOBEC mutation motif also contained nearly equal numbers of C to T and C to G events. Interestingly, spectra of scattered APOBEC motif mutations in APOBEC- hypermutated tumors were slightly, but statistically significantly, shifted towards C to T changes (Figure 2 and Figure S1). In general, the contribution of C to G changes in yeast models and in human cancers appear to increase, when long persistent ssDNA is expected to form.

**Figure 2.**
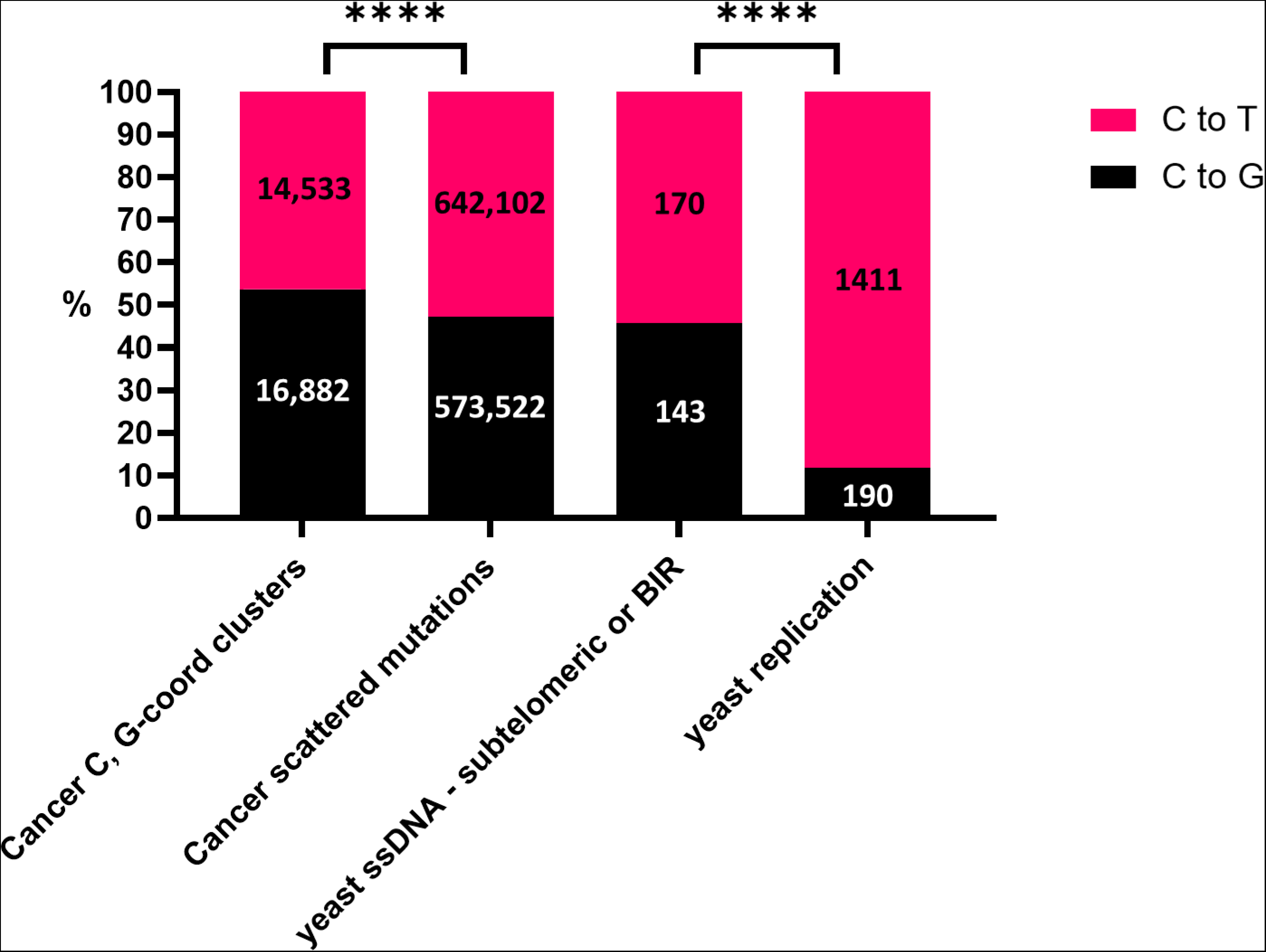
APOBEC-induced C to T and C to G mutagenesis in the data from PCAWG cancers and from yeast model studies. Yeast data include all C to T and C to G mutations. Counts for sections of yeast genome rendered single-stranded (subtelomeric 5’→3’ resection in uncapped telomeres or DSB-induced BIR (Chan *et al*. 2012; Chan *et al*. 2013; Hoopes *et al*. 2017; Elango *et al*. 2019)) were totaled into “yeast ssDNA” category. Counts for whole-genome sequenced yeast cultures studies (Taylor *et al*. 2013; Hoopes *et al*. 2016; Hoopes *et al*. 2017; Saini *et al*. 2017) were totaled into “yeast replication” category. Counts for individual yeast studies can be found in Supplementary Table 1. Cancer data include only C to T and C to G mutations found in tCw mutational motifs. Cancer data are totaled for all cancer types. Counts for each cancer type can be found in Supplementary Table 2, Figure S1 and in Supplementary Dataset 1. **** represents two-sided chi-squared P-value<0.0001

The universal feature of ssDNA formed in eukaryotes is that it readily binds to Replication Protein A (RPA) - multi-subunit protein complex required for DNA replication and for DSB repair (Chen and Wold 2014). Studies have shown that RPA binding can impede APOBEC cytidine deamination *in vitro* (Lada *et al*. 2011; Brown *et al*. 2021; Wong *et al*. 2021) and reduce APOBEC mutagenesis in yeast (Hoopes *et al*. 2016). Long persistent ssDNA resulting in the formation of APOBEC mutation clusters may have a lower fraction bound by RPA at any given moment, because of RPA depletion or because of continuously ongoing exchange between ssDNA-bound RPA and RPA in solution (Toledo *et al*. 2013; Chen and Wold 2014; Toledo *et al*. 2017). In order to explain the variations in the APOBEC mutation spectra in different genomic contexts (Figure 2), we propose that RPA may not only shield cytosines in ssDNA from APOBEC but can also shield APOBEC-induced uracils from uracil DNA glycosylase, thereby reducing a chance of creating AP sites (pathway (b1) on Figure 1) and consequently reducing a chance of C to G (and C to A) mutations. Another way for RPA to alter mutation spectrum of APOBEC-induced mutations could be via modifying choices of bases inserted by TLS across AP-site. Several lines of evidence suggested that RPA can play a role not only in DNA replication and repair but also in TLS by regulating PCNA sliding over ssDNA or/and by recruiting Rad6/Rad18 PCNA monoubiquitination essential for TLS (Hedglin and Benkovic 2017b; Hedglin and Benkovic 2017a; Hedglin *et al*. 2019). It may turn out that RPA modulation of TLS affects the TLS choice of inserting one of three possible bases across AP-site (step (b1.1) on Figure 1), thereby modulating spectrum of base substitutions in concert with ssDNA accessibility to RPA. Altogether, the RPA ssDNA binding and/or its’ interference with TLS may affect choice between C to T and C to G APOBEC mutagenesis. Therefore, we have explored the effects of a RPA hypomorph allele in the presence and in the absence of TLS capacity on the spectrum of APOBEC-induced mutations in a yeast *Saccharomyces cerevisiae* model. We present here results favoring the hypothesis that during DNA replication RPA shields uracils in ssDNA formed by APOBEC cytidine deamination from uracil DNA glycosylase Ung1, thereby allowing the direct copying of uracils by replicative DNA polymerases. Unexpectedly, we also found that for uracils shielded from Ung1 by wild-type RPA the mutagenic outcome is reduced by wild-type Pol zeta.

## MATERIALS AND METHODS

### Mutation reporter for accessing mutagenesis by APOBEC cytidine deaminases

The reporter design and rationale were described in (Figure 3) and in (Roberts *et al*. 2012; Hoopes *et al*. 2016). An additional single-base substitution reporter *ura3-29* (Shcherbakova and Pavlov 1996; Elango *et al*. 2019) contains an A:T to G:C mutation. This construct allows selection of the range of forward mutations inactivating *CAN1* gene by canavanine-resistance (Can-R). *CAN1* reporter was used to compare mutagenic effects of an APOBEC enzyme in strains of different genotypes. The reporter also enables selection of mutations in the *ura3-29* mutant cytosine base to one of the three possible bases, A, T, or G, because each of these substitutions and no other change in a yest genome can result in reversions of a strain to Ura^+^. Importantly, mutant cytosine is located within the trinucleotide context tCt (mutated base capitalized) preferred by most of APOBEC3 cytosine deaminases. The reporter cassette was placed on one of the two sides around the strong replication origin ARS216 thereby allowing the *ura3-29* cytosine to be mostly present in either lagging or in the leading strand template (left and right positions, respectively).

**Figure 3.**
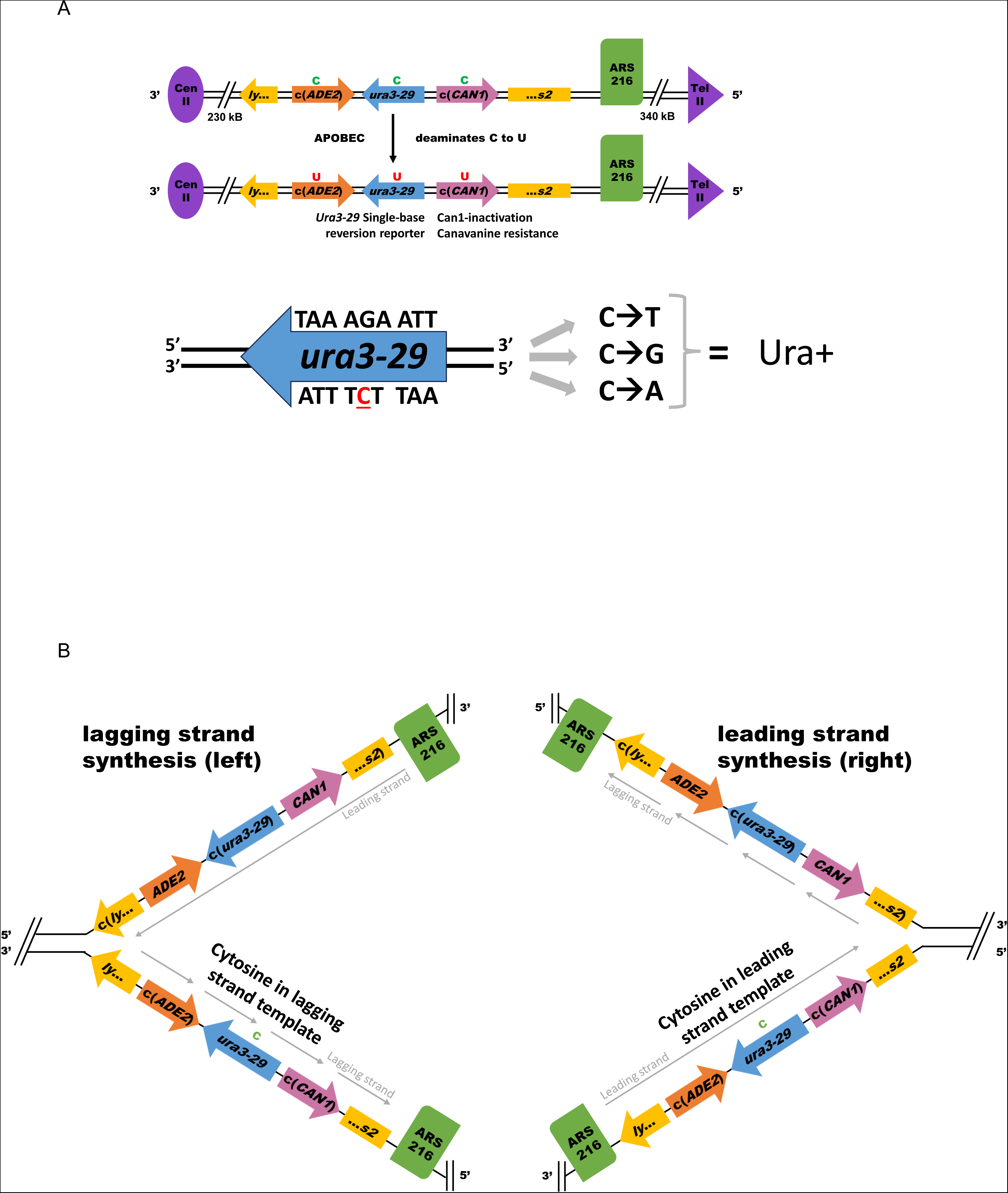
Yeast strains with *ura3-29* single-base mutation reporter for selection of three possible base substitutions resulting from TLS across AP-site. Details of all constructs of mid-chromosome triple-gene reporter were described in (Roberts *et al*. 2012; Hoopes *et al*. 2016) and in Materials and Methods. A. *ura3-29* mutation was placed within the triple-gene (*ADE2*, *URA3*, *CAN1*) mutation reporter, which was inserted into the LYS2 native ORF. Each of the three genes were deleted in their native positions. In addition to single base *ura3-29* reversion, *CAN1* gene allows to select forward mutations in the entire 1.6 kB ORF. **B.** Two orientations of triple-gene reporter around strong replication origin ARS216 place the mutant cytosine of the *ura3-29* in the lagging (“Left” construct; native position of LYS2 as in ySR128) or in the leading (“Right” construct originating from ySR366) strand template. Strain’s complete genotypes are listed in Supplementary Table 4.

### Yeast strains

The *Saccharomyces cerevisiae* yeast strains of CG379 genetic background used in this study were derivatives of the ySR128 (*MAT alpha ura3Δ can1Δ ade2Δ leu2-3,112 trp1-289 lys2::ADE2-URA3-CAN1*) in which triple-gene *ADE2-URA3-CAN1* was inserted into LYS2 in its normal chromosome II location to the left of ARS216 or of ySR366 (*MAT a ura3Δ can1Δ ade2Δ leu2-3,112 trp1-289 lys2Δ Chr.II 488694::lys2::ADE2-URA3-CAN1*) in which the entire ADE2-URA3-CAN1 cassette was moved on the left side of strong replication origin ARS216 (Figure 3 and (Roberts *et al*. 2012; Hoopes *et al*. 2016)). Standard methods were used to handle, grow, cross and dissect tetrads of yeast strains were used (Sherman *et al*. 1981). Introducing and PCR verification of the mutant *rfa1-t33* allele was described in (Hoopes *et al*. 2016). Single-base substitution *ura3-29* was introduced by integration and pop-out as described in (Shcherbakova and Pavlov 1996). Deletions of *UNG1* and *REV3* were generated by replacing their ORFs with natMX3 cassette, conferring resistance to nourseothricin or with the kanMX4 cassette, conferring resistance to geneticin, respectively (Goldstein and McCusker 1999). Combination of different genetic defects was done by mating type switching and crossing isogenic strains followed by tetrad dissection (Hoopes *et al*. 2016). Haploid progeny with a desired combination of generic defects was identified by phenotype and by PCR -verification. Yeast strains used in this work and their genotypes are listed in Supplementary Table 4.

### APOBEC plasmids

Vectors for the codon optimized expression of A3A, A3B, and A3G in yeast (i.e. pSR435, pSR440, and pCM-252-A3G, respectively) and the corresponding empty vector control (pySR419) were previously described (Chan *et al*. 2012; Hoopes *et al*. 2016). A1 (pSR433), A3C (pSR469), A3DE (pSR470), A3F (pSR471), and A3H (pSR472) yeast expression vectors were created by digesting gene-blocks containing codon optimized cDNAs (ordered from DNA2.0) with StuI and ClaI and ligating the resulting restriction fragment into the corresponding restriction sites of pySR419. Accurate cloning of the respective cDNAs was confirmed by Sanger sequencing of the inserted cDNA and DNA flanking the insertion site. Plasmid sequences in GenBank format are provided in Supplementary Dataset 3.

### Determining APOBEC-induced mutation rates

Yeast strains carrying an APOBEC plasmid or empty vector (EV) control plasmid were taken from genotype and phenotype-verified stocks (20% glycerol-80°C), patched on YPDA supplemented with 300 mg/L hygromycin (HYG) and grown for 2-3 days at 23 °C. These yeast strains were then streaked for single colonies on YPDA supplemented with 300 mg/L HYG with an addition of 20 mg/L doxycycline hyclate (DOX) to facilitate expression of an APOBEC ORF for 3 days at 30°C and to allow for large single colonies to form. The entire colony was then picked using a toothpick or pipette tip and suspended in 300uL of ddH_2_0 in a centrifuge tubes (dilution tube 0) followed by ten-fold serial dilutions. 100uL of 10,000x and 100,000x dilutions were plated onto COM media and grown for three days at 23°C to determine numbers of viable cells in a suspension. Non-diluted and 10x diluted suspensions were plated onto COM without arginine supplemented with 60 mg/L of L-canavanine or onto complete media lacking uracil and grown for 7 days at 23 °C. After the period of growth, the plates were stored at 4°C and the colonies were counted using the automated colony counter Protocol3. Mutation rates were calculated as described in (Saini *et al*. 2017).

### Collecting independent APOBEC-induced and spontaneous Ura+ revertants for determining spectra of base substitutions in *ura3-29* site and across the yeast genome

Independent Ura+ revertants were collected from small independent yeast culture grown on solid media as described in ((Jin *et al*. 2003) and references therein). Briefly small, about 1 uL, drops containing from 10^3^ to 10^4^ yeast cells were picked from yeast 10^5^ to 10^6^ cells/mL suspension with 121-pin multiprong device to solid YPDA media supplemented with HYG (300 mg/L) and 20 mg/L DOX for plasmid selection and for APOBEC induction, respectively, or directly to COM media without uracil to confirm that there was no or very little number of preexisting Ura+ mutants in the initial suspension. YPDA +HYG +DOX plates were incubated for 2-3 days at 30°C and then replica plated to COM media without uracil. By the time of replica plating each small prong imprint contained 2-5×10^6^ cells, so vast majority of Ura+ revertants on each prong occurred during growth of initially plated cells and therefore were independent from revertants on other prongs. One Ura+ revertant was picked from a single prong and streaked for single colonies. URA3 ORF was PCR-amplified, and Sanger sequenced or used for whole-genome sequencing. Primers used to amplify and sequence URA3 ORF from revertants are listed in Supplementary Table 4.

### Whole genome sequencing, mutation calling and mutation spectra analyses

Genomic DNA preparation, library preparation and Illumina sequencing and mutation calling was performed as described in (Hudson *et al*. 2023) with the exception that Illumina reads were mapped against yeast reference ySR128 (Roberts *et al*. 2012). Densities of APOBEC-induced mutations were presented as an average of mutation counts per genome for a group of APOBEC-expressing strain with the same genotype minus average density in the group of strains with the same genotype carrying the empty vector control plasmid. If the difference was negative, density of induced mutations was considered as zero. The average densities of induced mutations were used only for illustration purposes only. Set of whole-genome mutation counts in individual samples of the same genotypes were compared with the set of mutation counts in the group of another genotype using two-sided Mann-Whitney test. Mutation spectra were compared by two-sided chi-square of by Fisher’s exact test.

## RESULTS

### Increased APOBEC mutagenesis in yeast strains with hypomorph RPA allele *rfa1-t33*

As presented in the Introduction, we proposed that the increased fraction of APOBEC induced C→G changes in persistent long ssDNA as compared with ssDNA formed during normal replication could be explained by different access of Replication Protein A (RPA) to ssDNA formed in different genomic contexts. We employed *rfa1-t33* (S373P), a hypomorphic allele of the yeast RPA large subunit *RFA1*, to assess this hypothesis. The *rfa1-t33* was previously shown to reduce RPA DNA binding to ssDNA and to increase mutagenesis by APOBEC3A (A3A) and APOBEC3B (A3B) (Deng *et al*. 2014; Deng *et al*. 2015; Hoopes *et al*. 2016; Ruff *et al*. 2016). In extension of that finding we assessed effects of *rfa1-t33* on mutagenesis in *CAN1* gene by all human APOBECs for which deaminase activity has been reported (Refsland and Harris 2013; Salter *et al*. 2016; Mertz *et al*. 2022) (Figure 4). Mutagenesis was assessed in *ung1Δ* strains lacking uracil DNA glycosylase, therefore allowing all (or most of all) uracils generated by an APOBEC enzyme to be fixed into mutations (see Figure 1). The hypomorph *rfa1-t33* caused increased *CAN1* mutation rates even in the absence of APOBEC expression (EV, empty vector control) which could be explained by higher exposure of ssDNA to uncontrolled base-damaging factors and/or by increase in gross chromosome rearrangements encompassing *CAN1* (Banerjee *et al*. 2008). All APOBECs except APOBEC1 (A1) caused statistically significant increases of *CAN1* mutation rates over the empty vector control in *RFA1-WT* background, however, the increases in APOBEC3DE (A3DE), APOBEC3F (A3F), and APOBEC3H (A3H) expressing strains were only moderate, within 2-fold (Figure S2A). The same three APOBECs, A3DE, A3F, and A3H, did not result in statistically significant increases over the empty vector control in the strains carrying the hypomorph *rfa1-t33* (Figure S2B). On the contrary the *rfa1-t33* strains carrying A1, A3A, A3B, A3C, and A3G, showed statistically significant increases over the empty vector (Figure S2B), indicating that the increase in mutation rates was due to APOBEC-induced mutagenesis. As anticipated based on prior studies (see Introduction), the comparison of *CAN1* mutation rates between *RFA1-WT* and *rfa1-t33* strains also revealed the increase in APOBEC-induced mutagenesis caused by a the hypomorph allele in yeast carrying A1, A3B, and A3C (Figure 4). The strongest increase in rfa1-t33 over RFA1-WT background was observed for A1. This agreed with biochemical experiments that showed the strongest inhibition of A1 deaminase activity by human RPA *in vitro* (Wong *et al*. 2021). The weakest *rfa1-t33*-associated increase was for A3B which has the highest APOBEC-induced mutagenesis in both, *RFA1-WT* and in *rfa1-t33* backgrounds. Moreover, two other highly active enzymes, A3A and A3G did not show difference between *RFA1-WT* and in *rfa1-t33.* This minimal or non-existent difference in mutagenesis with the highly active APOBECs can be explained by the excess of enzymatic activity over the available ssDNA substrate. Altogether, our results indicate that *rfa1-33* hypomorph can facilitate access of an APOBEC enzyme to ssDNA in our strain background and therefore can be used to test our hypothesis about differences in RPA-ssDNA interactions accounting for differences in the spectra of APOBEC-induced mutation spectra in different genomic contexts (Introduction and Figure 2). We chose for this purpose the yeast strains expressing A3A, because this enzyme showed robust mutagenesis in both *RFA1-WT* as well as in *rfa1-t33* strains (Figure 4). Also, A3A is the most potent source of APOBEC-induced scattered and clustered mutations in APOBEC- hypermutated human tumors (Chan *et al*. 2015; Petljak and Maciejowski 2020; Petljak *et al*. 2022).

**Figure 4.**
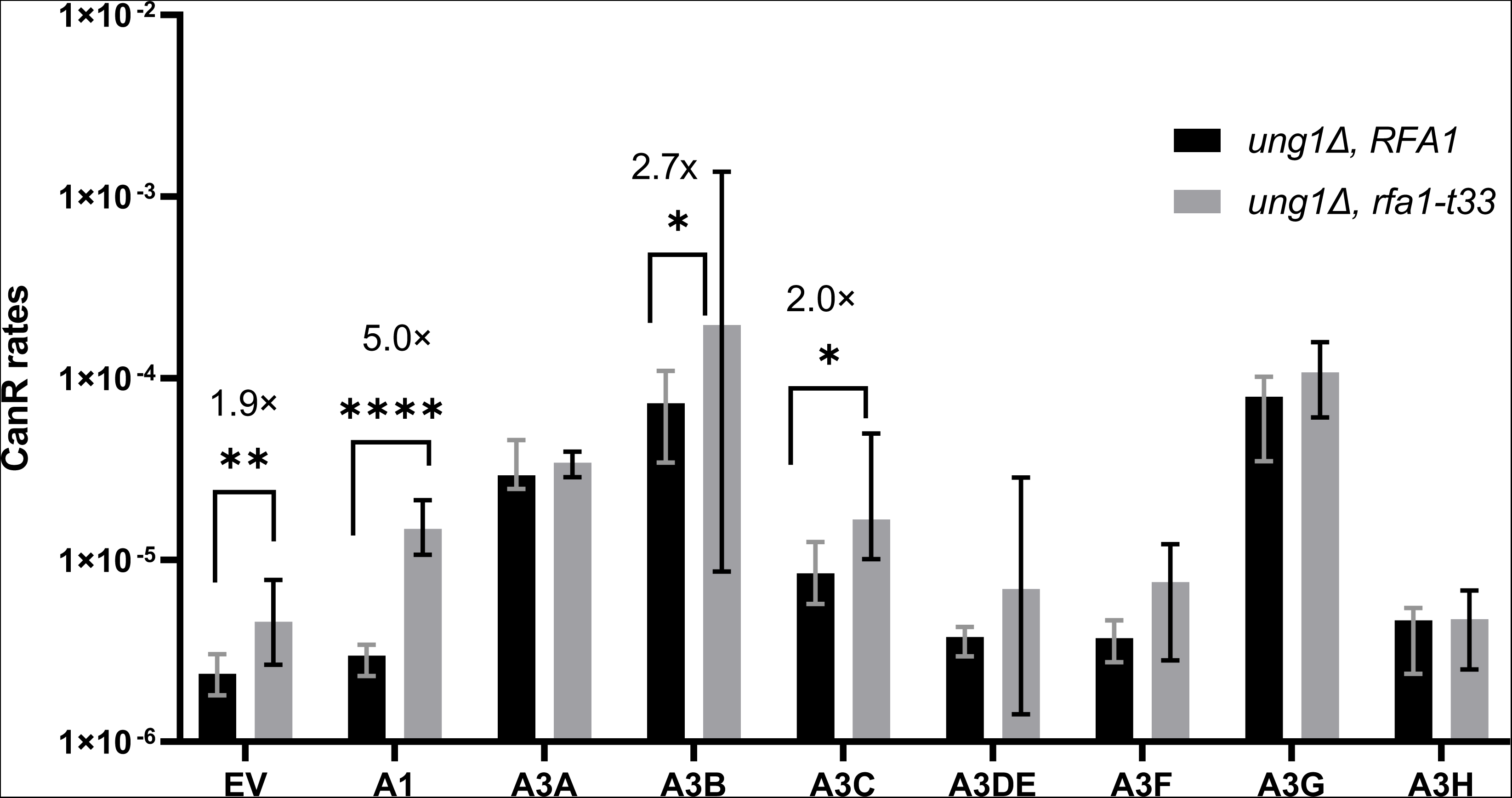
Stimulation of APOBEC mutagenesis in yeast strains carrying hypomorphic allele of RFA1. CanR mutation rates were measured in *RFA1* and *rfa1-t33* strains, all carrying a deletion of *ung1*. Shown are median values for mutation rates measured in 6-12 independent cultures and 95% confidence intervals for the medians. Statistically significant difference with P<0.05 of the two-tailed Mann-Whitney test is shown by brackets. Strains expressing A3B showed statistical significance only in one-tailed Mann-Whitney test. Source data and statistical analyses can be found in **Supplementary Table 4**.

### Uracil DNA glycosylase Ung1 deletion and *rfa1-t33* hypomorph can affect spectrum of base substitutions induced by APOBEC3A in *ura3-29* single-nucleotide reporter

As outlined in the Introduction, we hypothesized that the excess of C to G changes in APOBEC-induced mutation clusters as compared with mutations induced in the course of normal replication could be due to RPA shielding ssDNA in replicating cells from Ung or due to differences in choices of TLS across AP-sites in different genomic contexts. In order to explore these hypotheses, we compared mutation spectra induced APOBEC3A in glycosylase-proficient (*UNG1-WT*) replicating yeast between strains carrying *RFA1-WT* and *rfa1-t33* hypomorph allele in the presence and in the absence of TLS function (*REV3-WT* and *rev3Δ*, respectively). We also studied mutagenesis in *ung1Δ* strains, where no APOBEC-induced AP-sites were expected.

Mutagenesis was assessed with *ura3-2*9 single-base reporter allowing the determination of the mutation spectrum by sequencing *ura3-29* mutation site in the revertants. As expected, the Ura^+^ mutation rates in *ung1Δ* strains, where all mutations originated from mere copying of uracils, were generally higher than in *UNG1-WT*, where mutations can originate from uracils as well as from AP-sites (compare Figures 5A and 5B). In agreement with (Hoopes *et al*. 2016), there was a small but statistically significant increase in reversion rates of *ura3-29* positioned to the left of strong replication origin with the mutant cytosine would be mostly present in the lagging strand template. This bias was observed in wild-type strains with or without Ung1 as well as in *ung1Δ RFA1-WT rev3Δ* strains. Because of orientation dependence of ura3-29 reversion rates, both orientations were included into analysis of the mutation spectrum. Since reversion rates in in *UNG1-WT rfa1-t33 rev3Δ* strains were very low and did not produce enough revertants, *ura3-29* mutation spectrum was assessed only in *REV3-WT* strains (Figure 6). Mutation spectra were very similar between two orientations (Left and Right sides of the replication origin) therefore they will be considered as biological repeats in spectra analyses. As expected, all changes in *ung1Δ* strains were C to T regardless of *RFA1* genotype (Figure 6A). On the contrary, the spectra in UNG1-WT strains contained C to T as well as C to G and C to A mutations (Figure 6B). Mutation spectra in *UNG1-WT RFA1-WT* were mostly C to T with a very small fraction of C to G. This was similar to previously published whole-genome and reporter-based spectra of APOBEC-induced mutations in replicating yeast (Figure 2). Strikingly, the spectrum in replicating yeast strains of *UNG1-WT rfa1-t33* genotype contained nearly equal amounts of C to G and C to T mutations also with small presence of C to A. Thus, hypomorph *rfa1-t33* allele shifted the spectrum of APOBEC-induced mutations in replicating yeast strains to resemble the spectra observed in mutation clusters formed in long persistent ssDNA in yeast as well as in human cancers (Figure 2).

**Figure 5.**
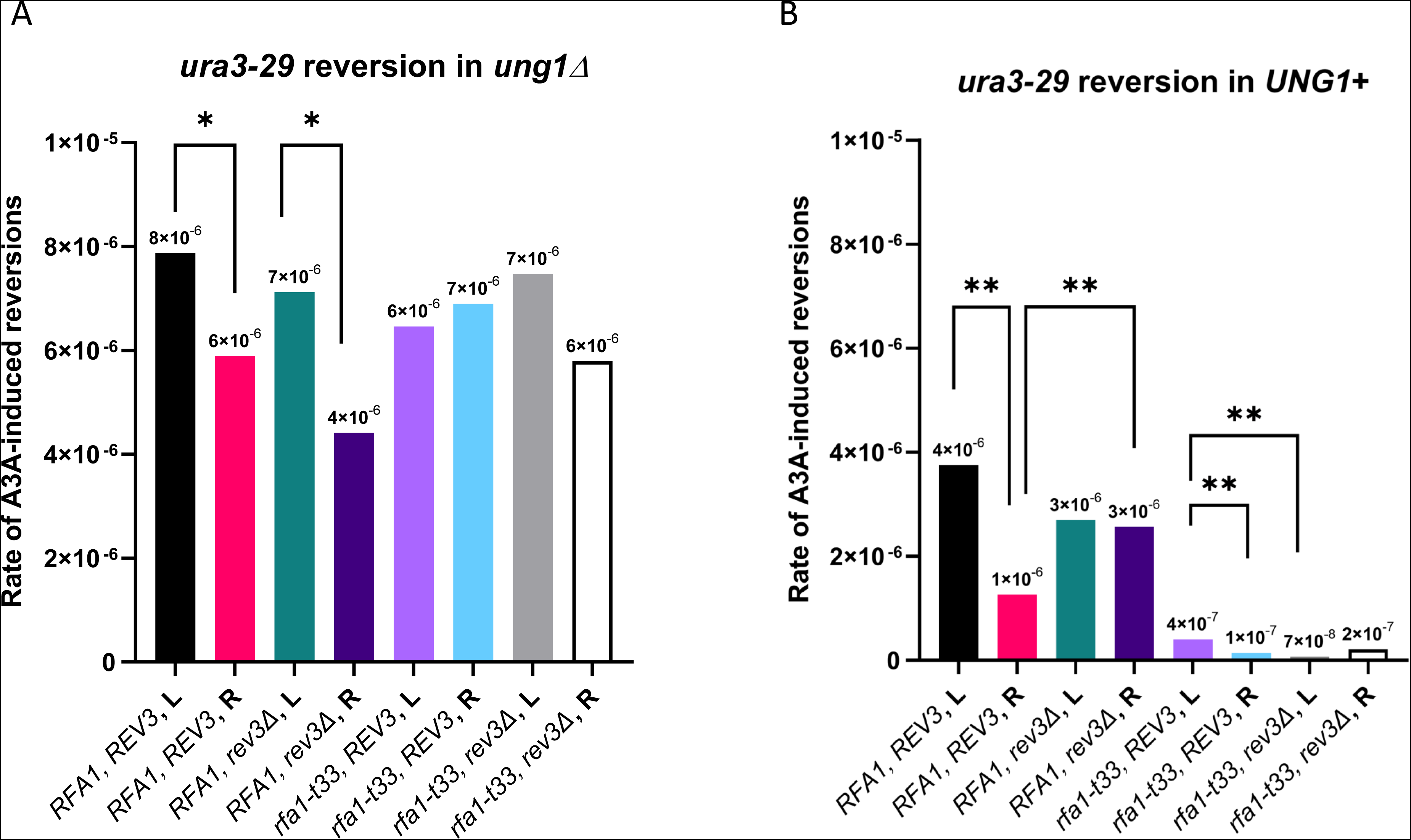
Effects of replication fork direction and strain genotype on rates of APOBEC3A- induced mutagenesis in *ura3-29* reporter. For all strains mutation rates were determined in 6 independent cultures. Rate of induced mutagenesis in each strain was calculated by subtracting median mutation rate in empty vector strain from a median mutation rate in A3A-expressing strain of the same genotype. Statistical comparison was performed by comparing sets of rates in a pair of strains by Mann-Whitney two-sided test. Pairs of strains showing statistically significant differences between mutation rates are connected with brackets. P-values are indicated as: * <0.05; ** < 0.005. **A, B** – *ura3-29* APOBEC3A-induced reversion rates in ung1- and in UNG1+ strains, respectively. Source data for panels **A** and **B** including details of statistical comparisons for P-values<0.05 can be found in Supplementary Table 5.

**Figure 6.**
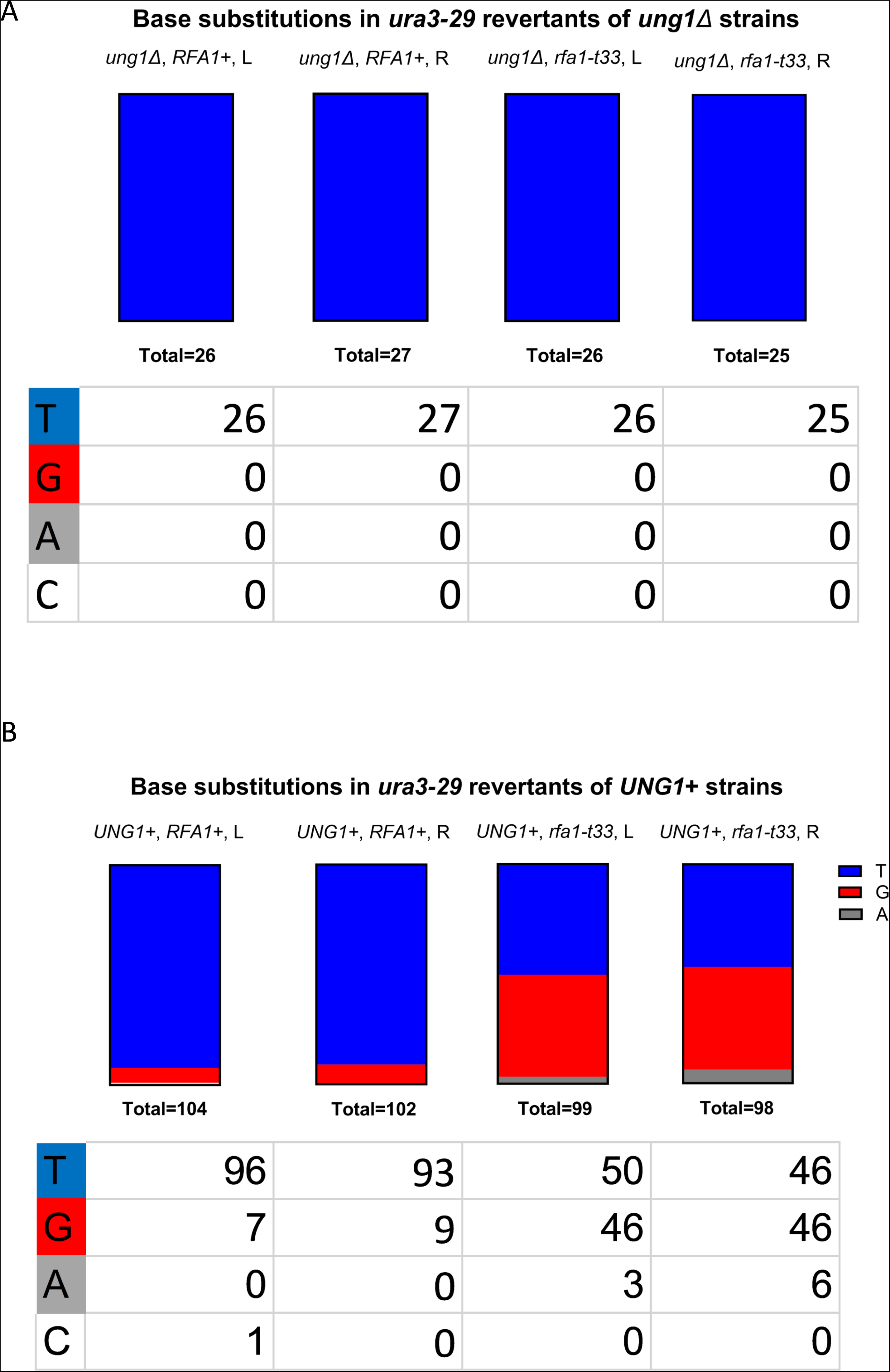
Base substitution spectra of APOBEC3A-induced ura3-29 changes in Ura^+^ reversion events. **A.** Mutation spectrum in *ung1Δ* strains. **B.** Mutation spectrum on UNG1-WT strains. Shown are percentages and numbers of base substitutions in ura3-29 mutation position of various genotypes. A single isolate without base substitution in ura3-29 position also included. For comparisons between spectra C to G and C to A changes were pulled to a single category. The only event with no changes in ura3-29 site was not included. Pairwise comparisons were performed using two-sided Fisher’s exact test. There were no statistically significant differences in spectra between strains of the same genotype carrying ura3-29 to the left (L) or to the right (R) of ARS216 (see Figure 3). Spectra in *RFA1-WT* strains showed strong difference from spectra in *rfa1-t33* strains (P<0.0001). Source data and complete outputs of statistical analyses of differences between spectra in different genotypes can be found in Supplementary Table 6.

### Hypomorph *rfa1-t33* allele increases probability of APOBEC-induced uracils being converted into AP-sites by Ung1 glycosylase

As outlined in the Introduction, the increase in C to G (and C to A) mutations of *ura3-29* in *rfa1-t33* strains could be explained by either increased probability of AP-site generation or by altering TLS preference to more frequently inserting cytosines across AP-sites. This question can be resolved by assessing mutation spectra in yeast *rev3Δ* lacking TLS capacity. However, *ura3-29* reversion rate in *UNG1-WT rfa1-t33 rev3Δ* strain was very low (Figure 5B). In order to assess the spectrum and frequency of APOBEC-induced mutations in all strains, including infrequently mutating *UNG1-WT rfa1-t33 rev3Δ*, we have sequenced genomes of 30-48 isolates of *UNG1-WT* strains belonging to each of the four genotypes in which we initially measured rates in *ura3-29* mutation reporter. Whole-genome sequencing allowed to build representative mutation catalogs for all strains, regardless of mutation rate. It also alleviated the possible impact of genomic context and of sequence composition in reporters. After subtracting empty vector background from mutation spectrum in all four genotypes we found that over 95% of remaining substitutions were in C:G pairs (Figure S3), which indicated that these are indeed A3A-induced mutations. Consistent with observations for *ura3-29* single-base substitution reporter in *UNG1-WT* strains (Figure 6A), whole-genome mutation spectra of *UNG1-WT rfa1-t33* contained more C to G mutations than the spectrum of *UNG1-WT RFA1-WT* strain (Figure 7A, B). (We note that all genotypes contained a tiny fraction of C to A mutations).

**Figure 7.**
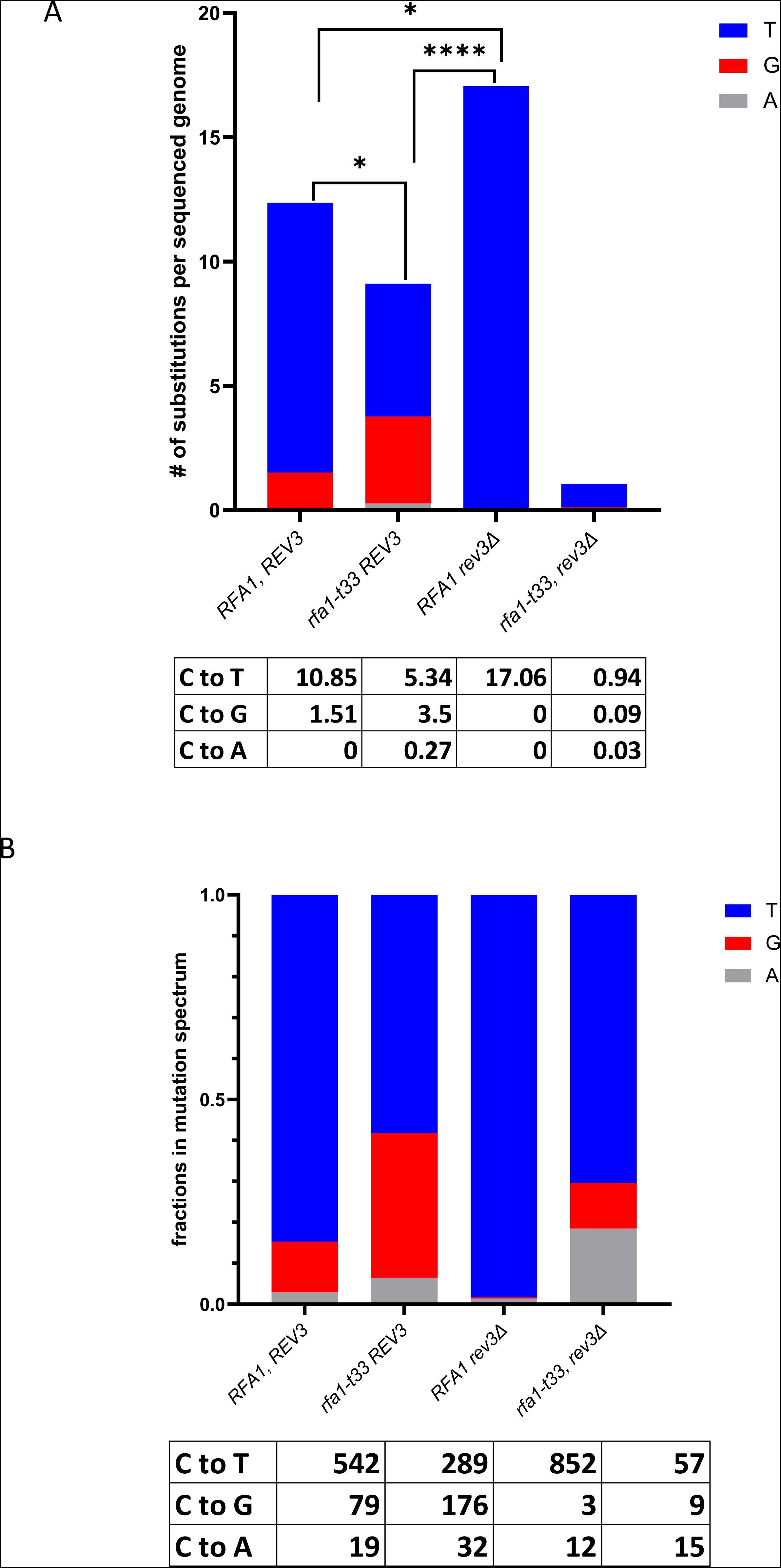
Whole-genome APOBEC3A-induced mutagenesis in yeast strains carrying wild-type allele of *UNG1*. Mutation calls can be found in Supplementary Dataset 2. **A.** Densities of APOBEC3A-unduced C:G base pair substitutions per sequenced genome (see Materials and Methods). Shown are substitutions in cytosines (C) of both DNA strands. Pairwise statistical comparisons between genotypes were performed by two-sided Mann-Whitney test comparing sets of C:G pair substitution counts in the sets of sequenced genomes of each genotype. All pairs with one exception produced P<0.05. For the comparison between sequenced isolates of *RFA1 REV3* and *rfa1-t33 REV3* strains, the Mann-Whitney test did not show statistically significance, however statistically significant differences of the mean values were confirmed by t-test which assumed that standard deviations of mutation calls were the same for the sets of sequenced isolates of these strains. Source data and complete outputs of statistical analyses of differences between different genotypes can be found in Supplementary Table 7. **B.** Total counts and fractions of total mutation counts for different substitutions in C:G base pairs of all genomes of a given genotype. All pairwise comparison between all pairs of genotypes by Chi-square showed strong differences (P<0.0001). Source data and complete outputs of statistical analyses of differences between different genotypes can be found in Supplementary Table 7.

While C to G (and C to A) mutations in UNG1-WT yeast can occur only from TLS of AP-sites generated by Ung excision of A3A-created uracils, C to T mutations can stem from either TLS of AP-sites or from error-free replication of uracils (Figure 1). When all branches of TLS across AP site were eliminated by *rev3Δ* in *rfa1-t33* background, densities of both C to G (and C to A) as well as C to T mutations were reduced more than 10-fold (Figure 7A). Such a severe reduction in densities of all types of APOBEC-induced substitutions suggested that bulk of A3A mutagenesis in *UNG1-WT rfa1-t33* was caused by AP-sites. There was smaller, albeit detectable presence of C to G (and C to A) mutations in *UNG1-WT RFA1-WT* strain, however the major fraction of APOBEC-induced mutations was due to C to T changes (Figure 7A, B). Elimination of TLS by *rev3Δ* in *RFA1-WT* strain left only C to T mutations in the spectrum but did not reduce mutation density (Figure 7A), therefore indicating that the majority of APOBEC-induced mutations in *RFA1-WT* replicating yeast resulted from copying of uracils that escaped Ung glycosylase processing into AP-site. Since most mutations in *rfa1-t33* had come from AP-sites, we conclude that the wild-type RPA can protect ssDNA of replicating yeast cells not only from APOBEC deaminases (Figure 4) but also from uracil DNA-glycosylase.

Unexpectedly, the density and fraction of C to T mutations increased in *UNG1-WT rev3Δ* strain as compared with *UNG1-WT REV3-WT* (Figure 7A). Since in *rev3Δ* all C to T mutations should originate from replication of APOBEC-induced uracils, the observed increase indicates that more uracils are formed and/or retained until replication. More studies are needed to understand the causes of the observed stimulation of C to T mutagenesis in the absence of Rev3 protein. Speculations about possible mechanisms are presented in Discussion.

## DISCUSSION

Our study revealed that in replicating yeast the contribution of two major base substitution types, C to T and C to G, into spectrum of APOBEC-induced mutations relies on capacity of RPA. Measurement of reporter-based and whole-genome mutagenesis in yeast carrying combinations of defects, hypomorph mutation in RPA large subunit *rfa1-t33*, uracil DNA glycosylase *UNG1*, and a deletion of a gene for the catalytic subunit of TLS polymerase Pol zeta (*rev3Δ*) indicated that the impeded functionality of RPA increases a chance to generate AP-sites from uracils formed by APOBEC cytosine deaminase in transient ssDNA (path (b1) on Figure 1). Replication across AP-sites by TLS can generate C to G as well as C to T mutations. Our results suggest a testable hypothesis about RPA counteracting Ung1 activity on ssDNA. Such a counteraction to Ung1 agrees with previously established inhibition of APOBEC cytosine deamination by RPA (Figure 4, and Introduction). Normal RPA could shield uracils in ssDNA from Ung1 the same manner that it shields cytosines from APOBECs. Thus, with normal RPA, most of APOBEC-induced uracils would be directed to pathway (b2) depicted in Figure 1, where only C to T mutations can occur.

Prevalence of the C to T mutations over C to G in wild type yeast replication (published data summarized in Figure 2 and the data of this study (Figures 6, 7) suggest that the latter base substitutions comprised only a minor fraction because RPA is protecting APOBEC-induced uracils from Ung in short-lived ssDNA within replication forks. Unlike replication forks, long, multi-kilobase stretches of ssDNA formed by end-resection of uncapped telomeres or in the course of BIR may exist for a much longer time and thus may give a greater chance for Ung acting on APOBEC-induced uracils. In addition, there could be some depletion of RPA in cells forming long persistent ssDNA (Toledo *et al*. 2013; Chen and Wold 2014; Toledo *et al*. 2017), which would further enable Ung access to uracils. Altogether, enhanced Ung function in long persistent ssDNA formed by end resection or in BIR would explain, why in these locations APOBEC induces approximately equal amounts of C to T and C to G mutations (Figure 2).

Our work also revealed unexpected result of increased C to T mutagenesis in *RFA1-WT* replicating yeast carrying deletion *rev3Δ* of the yeast Pol zeta essential for all branches of yeast TLS. In the absence of Pol zeta, even if AP-sites are formed by Ung (Figure 1 pathway (b1)) they could result in a base substitution. Some AP-sites could result in a loss of the DNA molecule (pathway (c)) and thus, no mutations are produced. Alternatively, AP-sites could be channeled into either error-free BER via lesion bypass or into reannealing of R-loop (Figure 1, pathway (b1.2), which would not produce mutations as well. Thus, changes within pathways generating AP-sites cannot explain increase in numbers of C to T mutations in the absence of Pol zeta. The only way to generate C to T mutations in the absence of Rev3 would be through accurate replication of uracils (Figure 1, pathway (b→b2→b2.1→b2.1.1)). At the moment there are no mechanistic knowledge suggesting how this pathway can be facilitated in the absence of Rev3 TLS. In principle, wild-type Rev3 could enhance channeling of uracils into error-free lesion bypass (Figure 1 (b2.2)) thereby reducing a chance for C to T mutations via accurate replication of uracils (b→b2→b2 →b2.2.1). It is well known that uracils impede archaebacterial DNA polymerases (Greagg *et al*. 1999; Richardson *et al*. 2013). On the contrary, uracil is readily copied by prokaryotic as well as eukaryotic replicases *in vitro* (Wardle *et al*. 2007). It remains to determine whether there is still some *in vivo* impediment to yeast replicases caused by uracils in DNA template. In support of possible direct effects of uracils in eukaryotes, recent studies indicated that uracils can trigger DNA replication stress in cancer cell lines even after siRNA suppression of Ung activity (Saxena *et al*. 2024).

Long and persistent stretches of ssDNA formed in cancer cells are the substrate for APOBEC mutagenesis resulting in clusters of multiple mutations in cytosines. In agreement with mutation spectrum observed for long persistent ssDNA formed in yeast APOBEC-hypermutated clusters in cancers contained nearly equal numbers of C to T and C to G mutations (Figure 2 and Supplementary Figure S1). While an unknown fraction of scattered (non-clustered) APOBEC-induced mutations in APOBEC hypermutated tumors may also be generated in persistent ssDNA, another part of scattered mutations would be coming from short-lived ssDNA in the replication fork. Indeed, a fraction of APOBEC motif C to T substitutions among scattered mutations was greater than the C to T fraction within APOBEC-hypermutated clusters. We note that the fraction of C to G substitutions mutations within the scattered mutation group in APOBEC hypermutated clusters is still higher than in replicating yeast. It remains to establish if the high fraction of C to T mutations is due to better access of human uracil DNA glycosylase (hUdg) to short lived ssDNA in replication fork or to other differences between yeast and cancer cells. Regardless of specific reasons, our work indicates that the ratio between C to T and C to G APOBEC-induced mutations can serve as a potential readout of the interplay between RPA protection of ssDNA and uracil DNA glycosylase action on uracils in ssDNA. The relative differences between C to T and C to G ratios in yeast and scattered replication associated APOBEC-induced mutations in human cancers highlights key differences in the dynamics of replication in these systems. The higher amounts of C to G substitutions in human cancers could indicate that the human RPA complex has a lower strength of ssDNA binding compared to the yeast complex. Alternatively, greater C to G could indicate the extent of RPA exhaustion and replication stress conditions present within tumor cells.

## DATA AVAILABILITY

All numeric and mutation call data necessary for confirming conclusions of this paper are presented in the text, in main and in supplementary display items, in supplementary tables and in supplementary datasets. Supplementary tables also contain numeric source data for each graph on display items. Illumina sequence reads were submitted to NCBI Sequence Read Archive under BioProject PRJNA1128109

## ACKNOWLEDGEMENTS

We thank Drs. Natalya P. Degtyareva and Rajula Elango for advice on the manuscript. This work was supported by US National Institutes of Health Intramural Research Program Project Z1AES103266 to D.A.G.; National Institutes of Health grants R01 CA218112 and R01 CA269784 (to S.A.R.).

## Supplementary Items

**Figure S1.**
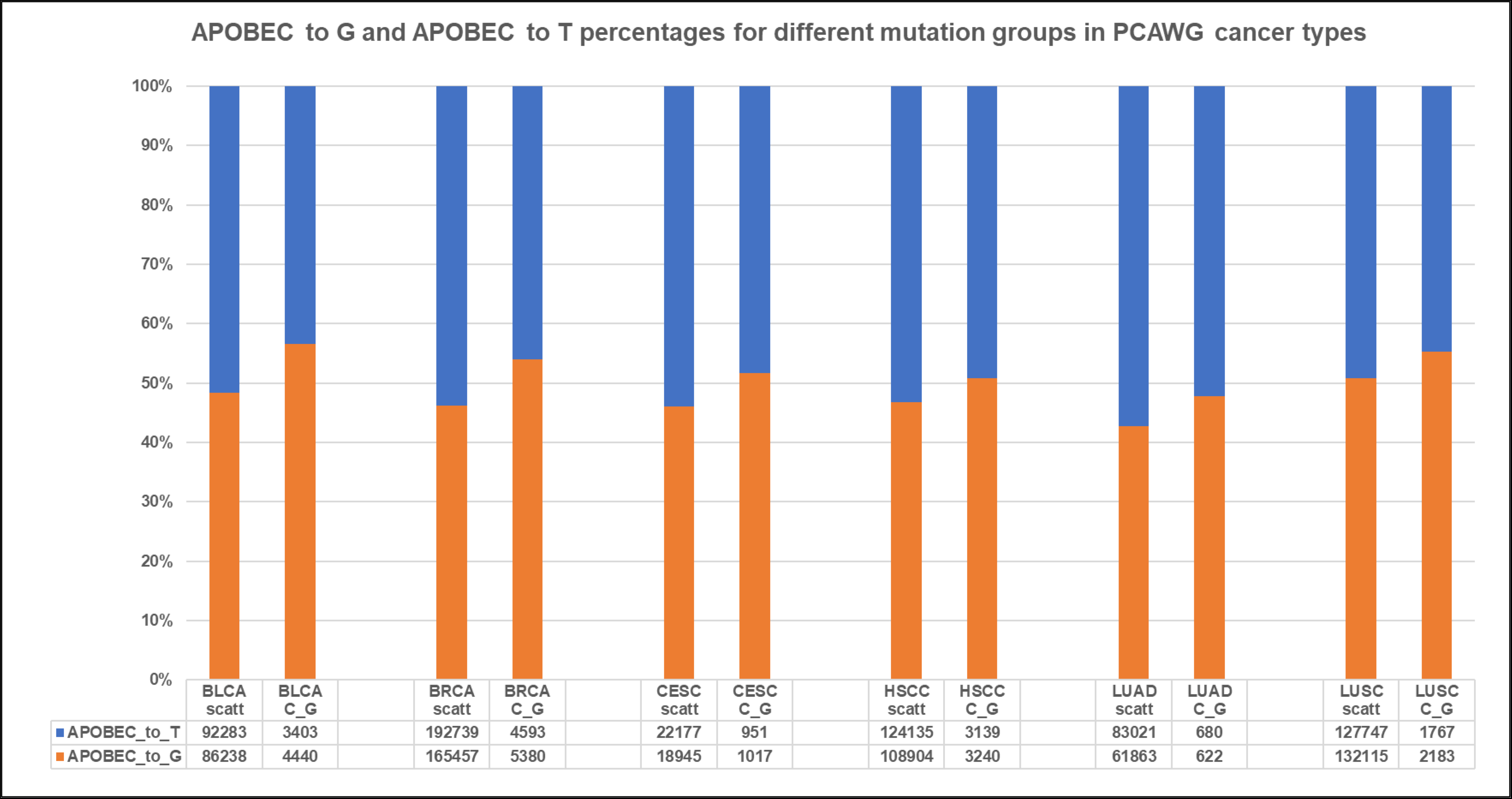
C to T and C to G mutations in APOBEC-hypermutated cancers. Shown are counts of mutation in APOBEC motifs tCw to T and tCw to G. Source data for this figure can be found in Supplementary Table 2. Cancer mutation catalogues and APOBEC analyses results can be found in PCAWG project data repository (Consortium 2020). Relevant summaries can be found in Supplementary Dataset 1. All mutation counts were taken from a row “Total” in columns ‘APOBEC_to_T’ or ‘APOBEC_to_G’of a cancer type. Separately calculated were values for mutations in C- or G-strand-coordinated clust ers (files *_sum_G_C_clusters_fisher_Pcorr.txt in Suplementary Dataset 1) or for mutations not assigned to any cluster (scattered mutations; files * _sum_scattered_fisher_Pcorr.txt). Cancer types are abbreviated as in TCGA (https://gdc.cancer.gov/resources-tcga-users/tcga-code-tables/tcga-study-abbreviations).

**Figure S2.**
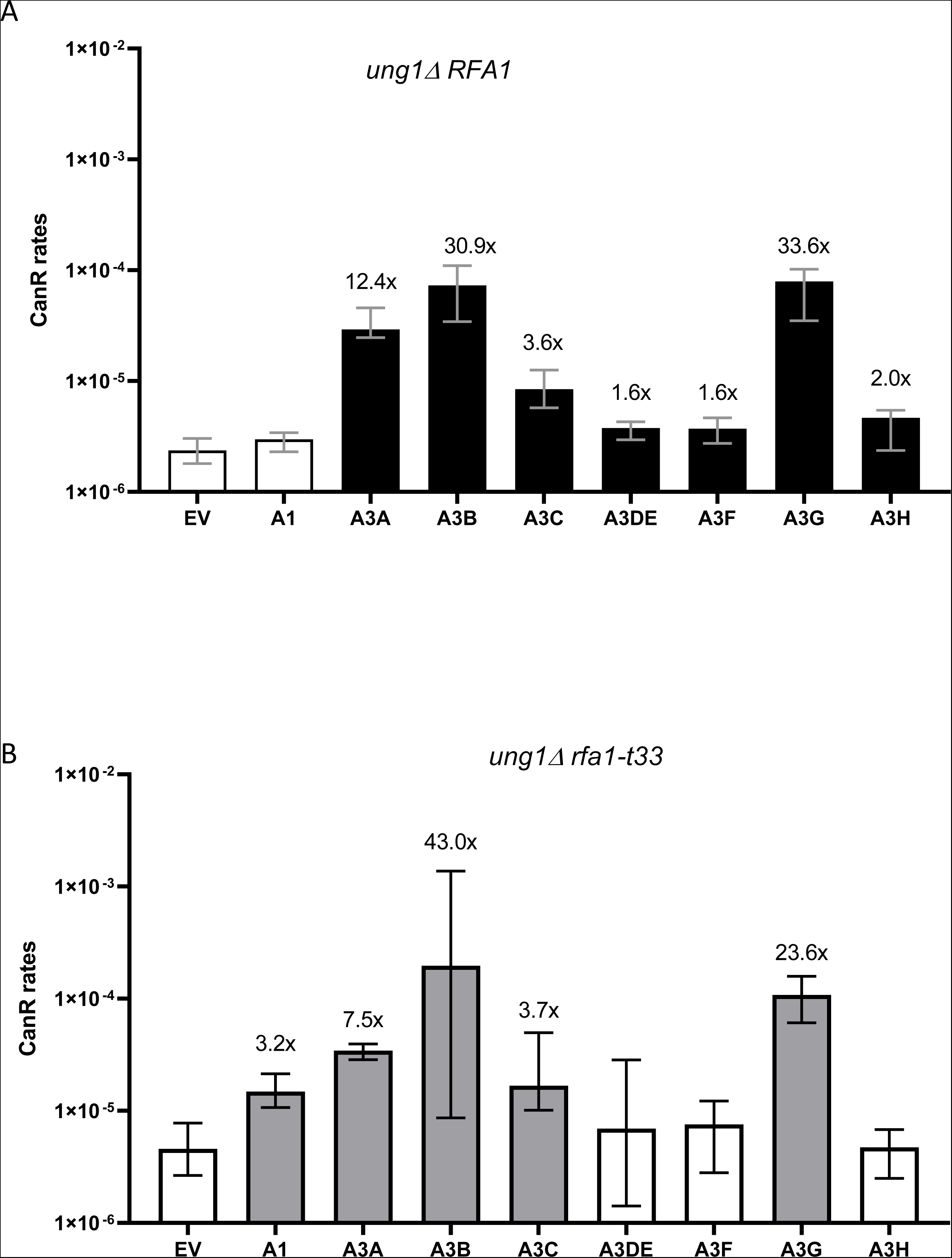
Comparisons of CanR mutation rates in yeast strains carrying vectors expressing different APOBEC enzymes with CanR rates in strains with empty vector. Filled bars correspond to APOBEC-expressing strains showing statistically significant (P<0.05) difference from the empty vector strain of the same genotype; numbers above filled bars indicate fold increases over empty vector. Source data and statistical analyses can be found in Supplementary Table 4. A. *ung1Δ RFA1-WT* strains. B. *ung1Δ rfa1-t33* strains.

**Figure S3.**
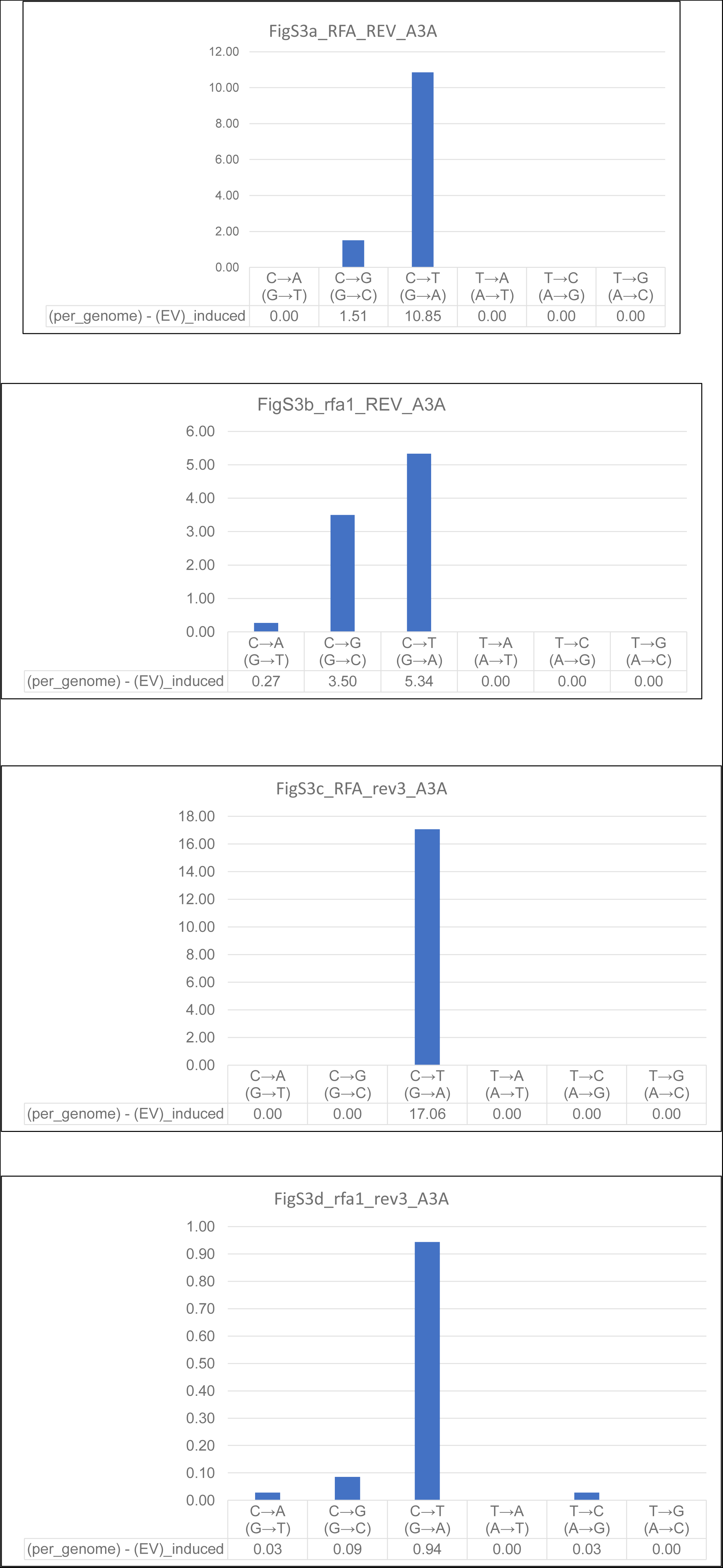
Calculated densities of APOBEC3A-unduced substitutions in C:G and A:T base pairs per sequenced genome.

## Notes

### Competing Interest Statement

The authors have declared no competing interest.

### Summary of Updates

This version of the manuscript has been revised to update the following - provide Supplementary Tables and Datasets

